# Fracture Mechanics of Human Blood Clots: Measurements of Toughness and Critical Length scales

**DOI:** 10.1101/2021.03.13.435277

**Authors:** Shiyu Liu, Guangyu Bao, Zhenwei Ma, Christian J. Kastrup, Jianyu Li

**Author notes:** Electronic Supplementary Information (ESI) available. See DOI: 10.1039/x0xx00000x.

## Abstract

Blood coagulates to plug vascular damage and stop bleeding, and thus the function of blood clots in hemostasis depends on their resistance against rupture (toughness). Despite the significance, fracture mechanics of blood clots remains largely unexplored, particularly the measurements of toughness and critical length scales governing clot fracture. Here, we study the fracture behavior of human whole blood clots and platelet-poor plasma clots. The fracture energy of whole blood clots and platelet-poor plasma clots determined using modified lap-shear method is 5.90±1.18 J/m^2^ and 0.96±0.90 J/m^2^, respectively. We find that the measured toughness is independent of the specimen geometry and loading conditions. These results reveal a significant contribution of blood cells to the clot fracture, as well as the dissipative length scale and nonlinear elastic length scale governing clot fracture.

## Introduction

Blood clots are hydrogels composed of fibrin and other proteins, infiltrated with platelets, red blood cells (RBCs) and other cells.^1^ They form at wounded blood vessels to stop bleeding, a process known as *hemostasis* (Fig. 1A). Blood clots plug and seal the blood vessel defects, where they are exposed to shear stress, tension and blood pressure from the local tissue environment (Fig. 1B).^1^ Pathological blood clots can appear within intact blood vessels, causing *thrombosis*.^2^ In both circumstances, the fracture of blood clots is detrimental and even fatal. The fracture of the blood clots at wound sites results in rebleeding (Fig. 1C).^3^ The fracture of an intravascular blood clot yields small pieces of thrombi that may disturb blood flow through the circulatory system and cause pulmonary embolism, heart attack or stroke.^4^ Thrombotic complications occur in many diseases; for instance, 31% of COVID-19 patients in intensive care units have venous and arterial thromboembolism.^5^ Understanding the fracture mechanics of blood clots may identify new mechanisms for controlling thrombosis and hemostasis.

**Fig. 1.**
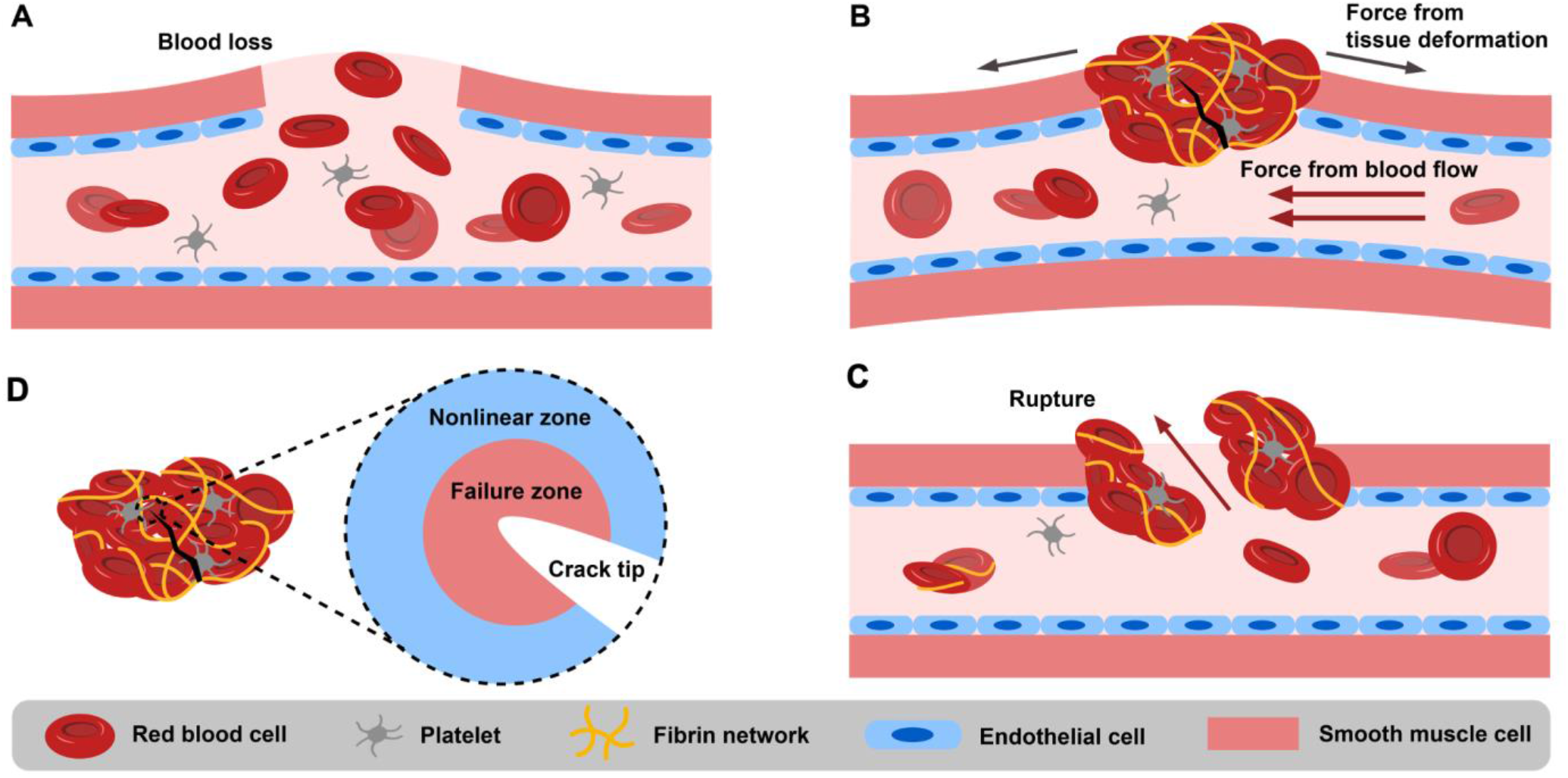
Schematics of clot formation and failure at a wound of a blood vessel. (A) Blood loss at a wound of a blood vessel. (B) A blood clot, formed to seal the defect, experiencing mechanical loading from the blood flow and the surrounding tissue, which might result in clot rupture. (C) Blood clots may rupture, causing rebleeding. (D) Schematic of the crack tip zone.

Despite the likely importance of fracture mechanics of blood clots in human health, it is not well understood compared to the fracture of hydrogels and many other soft materials.^6, 7^ Mechanical properties of blood clots such as stiffness, strength, and viscoelasticity are well characterized. Such properties, however, characterize blood clots that are intact and without cracks. Few works to date are reported on the toughness (*i.e.*, fracture energy required to extend a unit area of crack) of blood clots. Recently, the toughness of fibrin clots, derived from human blood plasma, was determined using single edge notched tension method and compared with finite element simulation.^8^ This work calls for further investigation on the human whole blood clots and the mechanisms underlying clot fracture.

As blood clots are fragile, wet and soft, characterizing the fracture properties of blood clots is non-trivial. Because blood clots are soft (shear modulus < 1 kPa) and deformable, this excludes using conventional fracture testing methods, such as three-point bending methods that are only applicable to rigid materials.^9^ Furthermore, because blood clots have low tensile strength <1 kPa,^8^ they are vulnerable to damage and even rupture during handling, transport and mounting, especially with a pre-crack introduced. It is problematic for many testing methods such as pure shear tests.^6^ Also, the wet nature of blood clots makes them difficult to grip or handle. Lastly, blood clots are not stable and there is a limited operational time window.

The fracture of blood clots involves mechanical, physical, and biochemical processes, given the structural and compositional complexity of blood clots.^1^ A running crack in a blood clot is expected to encounter complex stress/strain fields at the crack tip. There exists a failure zone where dissipation occurs and a nonlinear elastic zone where the material elements are deformed nonlinearly (Fig. 1D). The critical length scales to define the failure zone and nonlinear zone are known as the dissipative length scale and the nonlinear length scale, respectively.^10^ The crack tip zone is key to understand and model the blood clot fracture. For instance, the dissipative length reflects the critical flaw size, above which the clots sensitize to the crack.^11^ The critical length scales underling the clot fracture remains elusive.

In this work, we will apply fracture mechanics approaches to measure the fracture energy of human blood clots. We ask the following questions: what is the toughness of human blood clots and how does it depend on the composition and loading? To address them, we will first synthesize and characterize human whole blood clots and platelet-poor plasma clots using rheology. We will then combine modified lap-shear and double-cantilever beam methods to measure the toughness of the blood clots. To mimic aspects of tissue surfaces, we will employ rigid substrates coated with collagen casing, which can adhere with and support the blood clots for mechanical testing. The sensitivity of the measurements to the sample size and the loading condition will be examined. We will also estimate, for the first time, the dissipative and nonlinear elastic length scales, defining the crack tip field of blood clots. These findings will advance the understanding and modeling of blood clot rupture, which is an important mechanism in thrombosis and bleeding.

## Experimental Methods

### Materials

The human ethics protocol for this study has been approved by the Research Ethics Board at McGill University. Human whole blood (WB) and platelet-poor plasma (PPP) were purchased from ELEVATING SCIENCE®, BioIVT; the donors were unidentified. They were both pooled, drawn in citrate phosphate dextrose adenine (CPD-A) anticoagulant, and preserved in 1 mL aliquots. The human WB was kept at 4 ℃ and the PPP was frozen at −80 ℃. They were thawed at room temperature before use. Tissue factor was a recombinant thromboplastin reagent (Dade® Innovin® Reagent). Sample molds and cantilever beams were made of acrylic sheets after laser cutting. The acrylic and polyester backing films were from McMaster-Carr. Sodium chloride and calcium chloride dihydrate for preparing the recalcification solution were purchased from Sigma Aldrich.

### Blood clot formation

The blood clots were formed following a reported protocol.^12^ To initiate the clotting cascade, the recalcification solution containing NaCl and CaCl_2_ was mixed with the tissue factor at a volume ratio of 249:1. The mixture was then added into the blood samples (either WB or PPP), with a final volume ratio of the blood to the recalcification mixture solution at 3:1. While NaCl was fixed at 22.5 mM, the final concentration of CaCl_2_ was varied between 10 mM and 40 mM. Consequently, coagulation occurred as the clotting factors were activated and then cleaved and polymerized fibrinogen into a fibrin network. Immediately after recalcification, the blood clot was kept in a sealed Ziploc bag with saturated humidity inside to maintain hydration and the samples were incubated at 37 ℃ for 2 hours to allow complete coagulation and crosslinking prior to testing; the duration is sufficient according to the rheological measurement to be shown below.

### Mechanical testing

Mechanical tests were performed to characterize both the shear modulus and fracture energy of blood clots. The shear modulus was measured using a rheometer (Discovery HR-2, TA Instruments). The fracture energy was measured with the modified lap-shear method and modified double-cantilever beam method using a tensile machine (Instron 5966; 100-N load cell). Experimental details are provided below.

#### Rheological measurement

Rheology tests were performed to map the clotting kinetics of WB or PPP. Blood samples were recalcified as described above and deposited immediately onto the stage of a rheometer. The gap between the plate and the stage was set at 500 μm for a 160-μL blood sample. The temperature was set at 37 ℃ to mimic the physiological temperature. With the application of an oscillatory strain (1 %, 1 Hz) using a 20-mm parallel steel plate, the clotting kinetics was characterized by monitoring the storage (G’) and loss (G’’) moduli, which correspond to the elastic and viscous properties of the tested materials, respectively. The storage and loss moduli were recorded over time up to 3600 seconds. Coagulation time corresponded to the time when the storage modulus reached 90% of its plateau value. At least 4 replicates per condition were tested.

#### Modified lap-shear tests

Modified lap shear tests were performed to measure the fracture energy of the blood clots. This method has been widely applied to characterize hydrogels and adhesives.^13, 14^ Different from the conventional lap-shear tests, polyester films (100-μm thickness) were added as a rigid backing to constrain the axial tension of the specimen. The backing was glued to a collagen casing, yielding a tissue-mimicking surface. Such surfaces can adhere to the blood clots^15^, transmit the loading and guarantee cohesive failure, instead of adhesive failure. The recalcified blood (length 35 mm, width 15 mm, and thickness 0.1 mm) was placed between two collagen-coated backings, where a polyester film of 100-μm thickness was inserted to define the thickness of the specimen. An incision of 1-mm length was introduced as a pre-crack at the edge of the samples.

The specimens were loaded vertically at a displacement rate of 3 mm/min using the Instron machine. Following an established protocol^13^, the fracture energy, defined as the reduction of the potential energy associated with crack propagating per unit area, was calculated as the total work (i.e., the area under the force-displacement curve) divided by the area of the crack surface after rupture, Γ =*W_t_*/*A*. The shear strength was also calculated as the maximum force in the force-displacement curve divided by the initial area of the sample. The fracture surfaces were examined post-testing to confirm cohesive failure. At least 4 replicates per condition were tested.

#### Modified double cantilever beam tests

Modified double cantilever beam tests were also performed to measure the fracture energy following a prior work.^16^ Acrylic sheets were laser-cut and glued to produce rigid cantilever beams (60 mm x 10 mm x 6 mm) and coated with collagen casing to mimic tissue surfaces. The blood sample was recalcified and immediately placed between two collagen-coated acrylic sheets; its geometry (40 mm length, 10 mm width, and 0.1 mm thickness) was defined using the polyester film. The specimens were then incubated as described above and tested with the Instron machine. The displacement-controlled loading rate was 3 mm/min. Fracture energy was calculated as the total work (area under the force-displacement curve) divided by the area of the area of the crack surface after rupture, i.e., Γ =*W_t_*/*A*. At least 4 replicates per condition were tested.

#### Scanning electron microscopy imaging

The microstructures of WB and PPP were imaged using a field emission scanning electron microscope (SEM, F450, FEI). Before the SEM imaging, all the samples were dehydrated using a CO_2_ supercritical point dryer (CPD030, Leica) to preserve the original microstructure. The dehydrated samples were coated 4-nm Pt using a high-resolution sputter coater (ACE600, Leica) to increase surface conductivity.

### Statistical analysis

All analyses were conducted using Originlab Pro 2018. One-way analysis of variance (ANOVA) was used to compare differences between groups of experimental data. Both F-and P-values were calculated. All data are represented as a means ± 0.5SD, unless otherwise noted.

## Results and Discussion

### Rheological measurement of blood clots

We first characterized the blood clots with rheological tests to determine the shear modulus. Both WB and PPP were tested and compared for the role of blood cells in clot fracture. The blood clots were initiated with the mixture of recalcification solution and tissue factor (Fig. 2A). The addition of Ca^2+^ initiated the coagulation cascade and produced thrombin, while the tissue factor further amplified the generation of thrombin by an extrinsic pathway of coagulation, converting soluble fibrinogen into an insoluble cross-linked fibrin network.^17^ The kinetics of clot stiffening was mapped with rheological time-sweep tests. Figure 2B shows the storage modulus surpasses the loss modulus quickly and reaches a plateau, indicative of equilibrium.

**Fig. 2.**
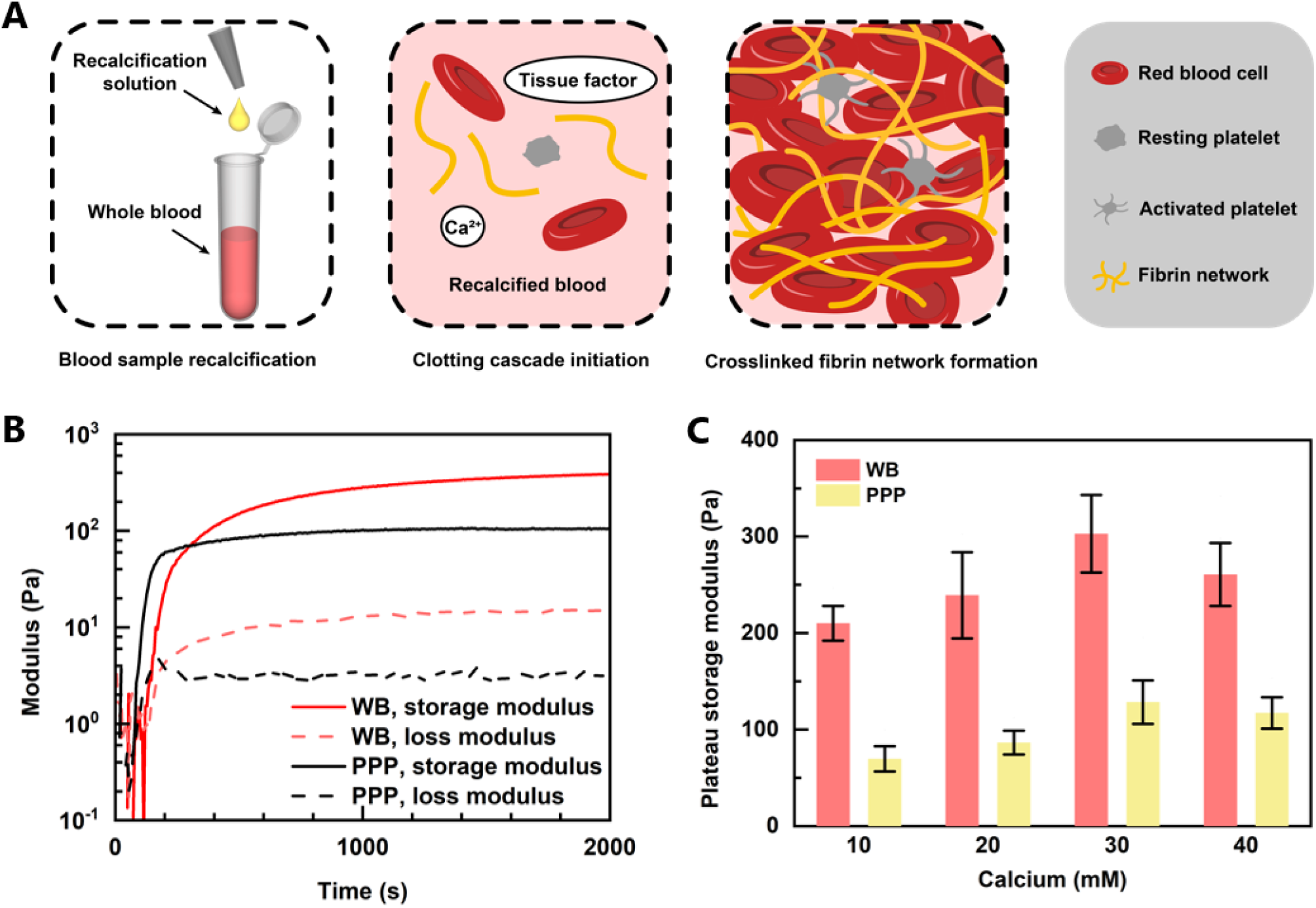
Rheological measurements of blood clots. (A) Schematics of blood clot sample preparation. (B) Gelation kinetics of clotting whole blood (WB) and platelet-poor plasma (PPP). (C) Plateau storage moduli of whole blood clots and platelet-poor plasma clots as a function of calcium concentrations. Sample size N=4; error bars, mean ± 0.5SD.

Blood clots stiffened with the presence of blood cells such as RBCs and platelets. To visualize the microstructure, we used SEM to image the microscopic morphology of the clots. The PPP >clot is a typical cross-linked fibrin network, while the WB clot is a tight aggregation of RBCs connected by the fibrin network and platelets (Fig. S1, ESI†). As the RBCs take up to 50% of the total blood volume, they affect the structure and mechanics of fibrin fibers and the resulting network.^18^ Fiber diameter increases upon RBC incorporation and RBCs also influence the viscoelastic properties of the clot.^19^ The incorporation of platelets leads to clot contraction and clot stiffening. ^20^

As calcium ions are reported to influence blood clot stiffness,^21,22^ we also measured the clot modulus with varying calcium concentrations. Both WB and PPP clots stiffen slightly as the calcium concentration increases from 10 mM to 30 mM (Fig. 2C) and the effect saturates at high calcium concentrations (30 mM and 40 mM). A similar trend was also observed on the change of coagulation time with increasing calcium concentration (SI fig. 2A, B). As a result, 30 mM final calcium concentration was used in the following study on both human WB and human PPP clots. The shear modulus of human WB clots is 302.92±80.42 Pa, and that of human PPP is 128.45±45.18 Pa. The measurements agree well with the literature,^22–24^ validating the blood clot formation. They will be used later for calculating the critical length scales underlying clot fracture.

### Lap-shear tests of blood clots

We next quantified the toughness of blood clots as fracture energy measured with modified lap shear tests. The testing method was chosen due to its applicability to soft fragile samples that might deform substantially due to gravity alone. Also, the test has been widely applied for characterizing blood clots, hydrogels, and other biomaterials.^13–15^ In this study, we used collagen-coated acrylic sheets, which adhere to and exert loading to blood clots. To examine the fracture behavior, we introduced a pre-crack in the middle of the sample and adjusted the alignment of the specimen and the loading direction to ensure the crack would propagate through the clot matrix (Fig. 3A). The rupture started when the force peaked. By examining the fracture surfaces of the specimen post-testing, we found sample residues on both sides of the backing, confirming cohesive rupture (Fig. 3B inset). Furthermore, SEM images reveal fibrin fibers pulled out at the crack tip (Fig.3A inset). From the force-displacement curves of the WB and PPP clot specimens, we determined the fracture energy of WB and PPP blood clots at 5.90±1.18 J/m^2^ and 0.96±0.90 J/m^2^, respectively (Fig. 3B, C). The shear strain rate was fixed 20/min in this study. It should be noted that the loading rate is expected to play a role as the blood clots are highly viscoelastic, calling for further investigation.^25^

**Fig. 3.**
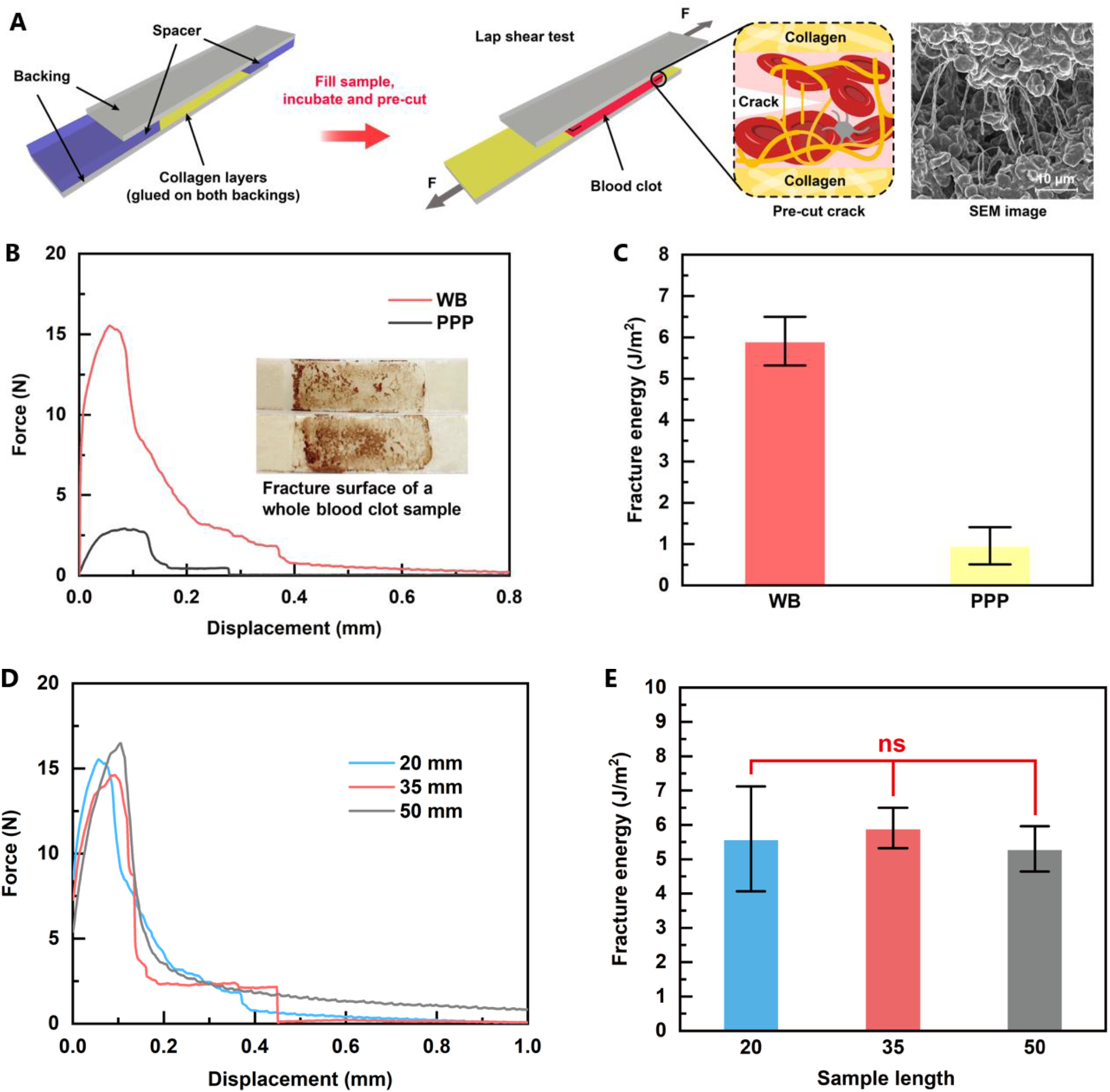
Fracture energy of blood clots measured by lap shear test were size independent. A) Schematics of lap shear test setup. (B) Force-displacement curves of lap shear tests for whole blood (WB) clots and platelet-poor plasma (PPP) clots; the inset shows the fracture surface of the WB specimen. (C) Fracture energy results of whole blood clots and platelet-poor plasma clots. Sample size N= 4 (D) Representative force-displacement curves of whole blood clot lap shear tests of different sample lengths. (E) Fracture energy of whole blood clots obtained from lap shear tests of different sample lengths. Sample size N=4; error bars, mean ± 0.5SD; NS: no significance, one-way analysis of variance (ANOVA), Tukey test.

The comparison between WB and PPP clots reveals the significant contribution of blood cells to the toughness of blood clots. RBCs and platelets influence the fibrin network, clot formation, maturation, stability, embolization, and fibrinolysis.^18,19^ The toughening effect contributed by blood cells is intriguing, and likely relates to viscoelastic and dissipative properties of blood cells (particularly, disrupting cell membranes of RBCs^26^) and their interactions with the fibrin network, which alter the fibrin network structure, individual fiber characteristics, and overall clot viscoelasticity.^19^

### Fracture energy measurements with varying loading

As fracture energy is expected to be invariant against the geometric factor and testing methods,^27^ we next measured toughness after varying the geometric and loading conditions.

First, we performed the lap-shear tests with varying lengths of WB clot samples (20-50 mm). The force-displacement curves and the measured toughness were compared (Fig. 3D, E). An ANOVA was conducted and identified no significant difference with F=0.0830 and P=0.9212.

We further changed the lap-shear method with a modified DCB method to test human PPP clots (Fig. 4A). In the modified DCB tests, collagen-coated acrylic sheets were use as rigid cantilever beams, which sandwiched and loaded the PPP clot. The fracture energy measured by modified DCB tests (0.95±0.15 J/m^2^) and by lap shear tests (0.96±0.90 J/m^2^) was consistent (Fig. 4B). In comparison, the DCB strength (5.59±0.36 kPa) differed from that of the lap shear tests (3.92±2.47 kPa) (Fig. 4C). This result demonstrated that fracture energy can be used to quantify the toughness of blood clots, which was independent of sample geometry and loading, as expected for a material property.

**Fig. 4.**
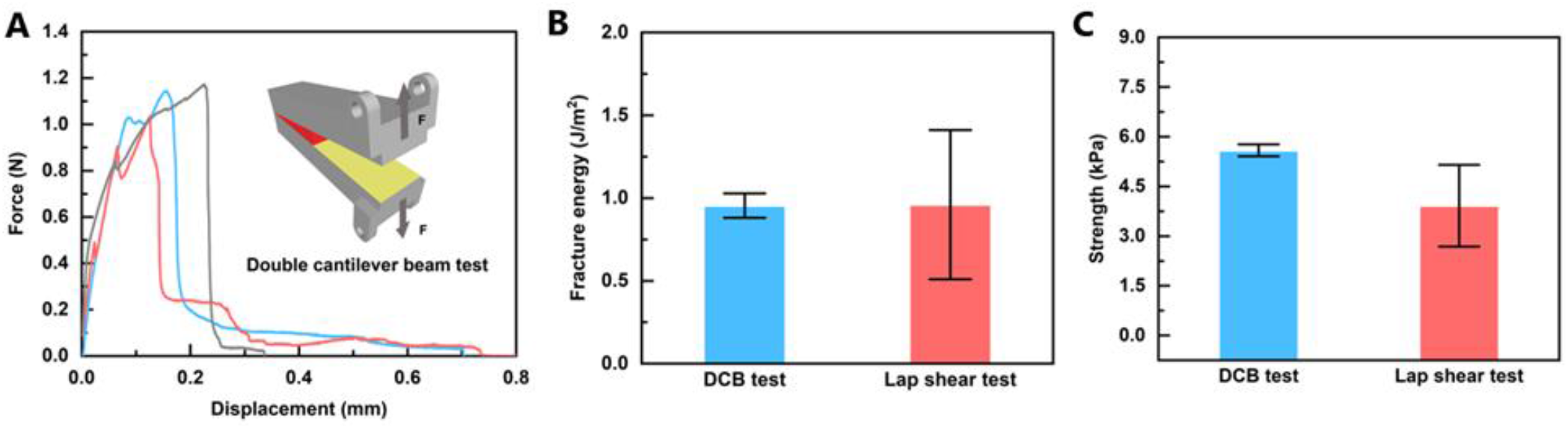
Lap shear test results were compared with double cantilever beam tests, and loading condition independence was found. (A) Schematics of double cantilever beam (DCB) testing, and representative force-displacement curves of DCB tests on PPP clots. (B) Fracture energy of PPP clots obtained from DCB tests and lap shear tests. (C) Strength of PPP clots measured by DCB tests and strength of PPP clots measured by lap shear tests. Strength is defined as the maximum force divided by the initial area of the sample section surface. Sample size N=4; error bars, mean ± 0.5SD.

### Length scales of blood clot fracture

We next estimated the critical length scales governing the fracture of blood clots: the dissipative length scale *ξ* and the nonlinear elastic length scale *l*.^11^ The former describes the size of the process zone near the crack tip during fracture and reflects the flaw sensitivity of a material, while the latter depicts the size of materials undergoing nonlinear deformation at the crack tip (as illustrated in Fig. 1C).^10^ The dissipative length scale was estimated by the ratio of the fracture energy to the work to rupture, *ξ* =Γ/*W_*_*. The work to rupture *W.* was characterized by loading specimens without a cut to rupture using the same setup (lap shear test in this study) and defined by the area under the stress-strain curve. The nonlinear elastic length scale, also known as the elasto-adhesive length, can be represented by the ratio of the fracture energy to the shear modulus, *l* =Γ/*E*. As can be seen, the length scales were on the order of 0.1-1 mm, reflective of large deformability and irreversibility (dissipation) of blood clots (Fig. 5A).

**Fig. 5.**
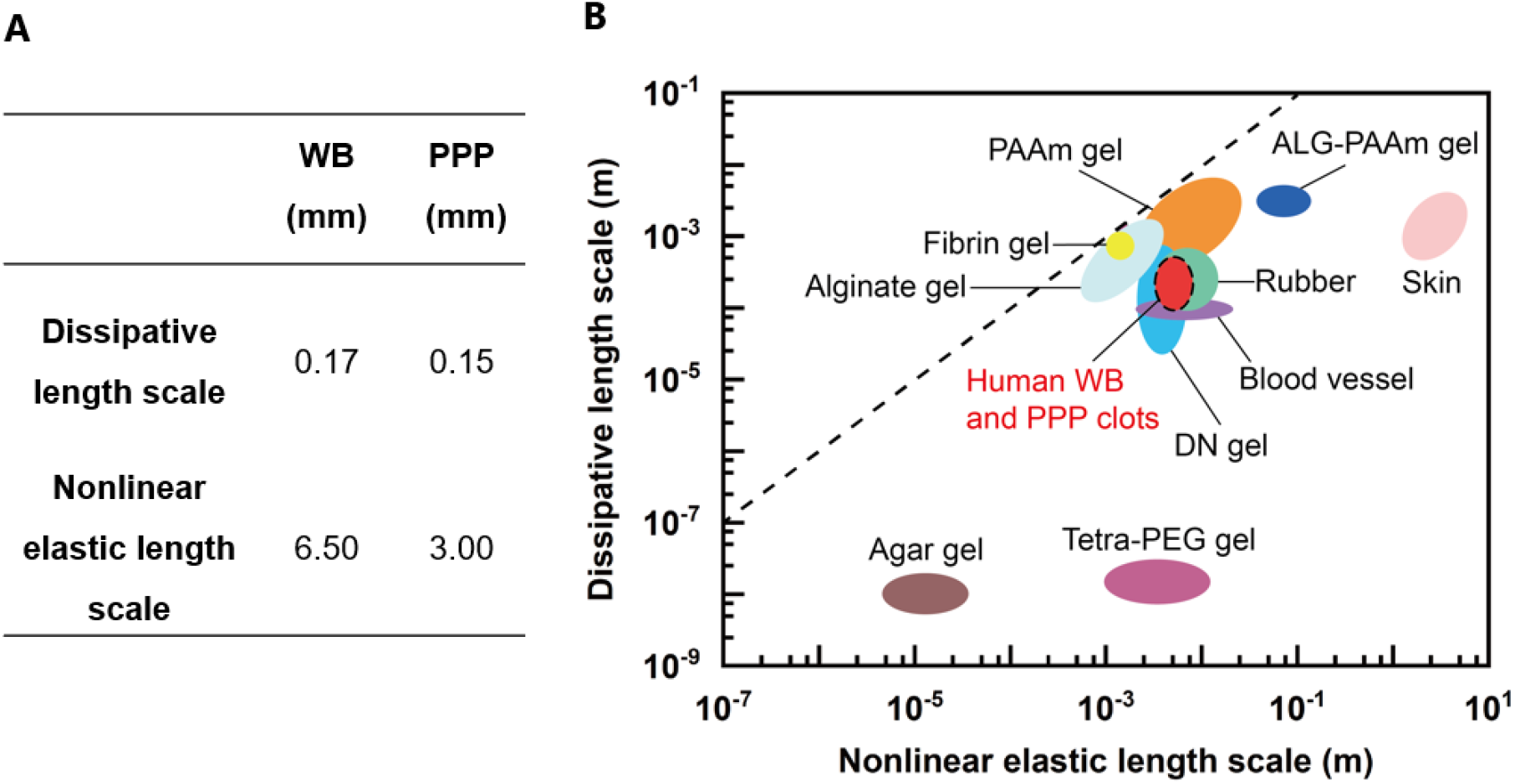
Dissipative and nonlinear elastic length scales of a variety of materials. (A) Rough quantified critical length scales of whole blood (WB) and platelet-poor plasma (PPP) clots measured in this work. As expected, the dissipative length scale is smaller than the nonlinear elastic length scale, since the blood clots are soft. (B) Fracture length-scale chart for soft and biological materials from reported work by others^8, 11, 28–34^. Dash line represents that the material’s dissipative length scale and nonlinear length scale are equal.

The two length scales provide insights for modeling the crack tip field. For example, when the crack size exceeds the dissipative length scale, the failure is governed by crack growth (fracture mechanics). Given a crack smaller than the dissipative length scale, the material is insensitive to cracking (flaw-insensitive) and thus follows the strength criteria instead.^10^ For highly deformable viscoelastic materials, the nonlinear length scale represents the area where the deformation is appreciably nonlinear and considerable energy dissipation occurs.^11^

To further compare the blood clots with other materials, we collected the dissipative and nonlinear elastic length scales for various materials. Figure 5B illustrates the values for human WB and PPP clots from this work, and those reported in literature for fibrin gel^8^, blood vessel^28, 29^, skin,^30^ natural rubber,^31^ alginate gels,^30^ double-network (DN) gels,^11, 32^ polyacrylamide (PAAm) gels,^30^ PAAm-alginate gels,^30^ agar gels,^33^ and tetra-arm polyethylene glycol (Tetra-PEG) gels^34^. Compared with the fibrin clot^8^, the WB clots have a smaller flaw sensitivity length scale (~0.2 mm) and larger nonlinear length scale (~7 mm), reflective of the fact that the WB clots contain large fraction of blood cells. Interestingly, the two length scales of human blood clots are comparable with that of double-network hydrogel and natural rubber. This finding implies that blood clots are relatively flaw insensitive and highly deformable, therefore have a great potential for fracture resistance.

The impressive length scales are associated with the unique structure and composition of blood clots. The porous fibrin network, as the main structural protein, endows high deformation and softness, while maintaining a sufficient permeability to allow effective enzymatic decomposition. Platelets attach to the fibrin fibers to stiffen and densify the clot, thereby providing a mechanically stable seal. The incorporation of soft and viscoelastic filler, RBCs, can dissipate the stress and mechanical energy when under strain. They are manifested as the hysteresis, Mullins effect and rate-dependent effect (viscoelastity) of blood clots under cyclic loadings.^25^ The detailed toughening mechanisms should be further investigated.

## Conclusions

To sum up, we investigated the fracture mechanics of human blood clots. We measured the fracture energy of human WB clots and PPP clots, as well as the critical length scales of blood clots near the crack tip. We characterized the fracture energy with modified lap-shear tests and found that it was insensitive to the sample size. The fracture energy measured with the lap-shear and DCB tests was in good agreement. We further calculated the dissipative length scale and the nonlinear elastic length scale of the blood clots. They provide valuable insights into the stress-strain field in the crack tip during blood clot rupture. Our study concludes that blood clots are highly deformable, dissipative and relatively flaw-insensitive. The underlying mechanisms are expected to link with the elastic nonlinearity and dissipation of the fibrin network and the blood cells, which calls for future investigations. This motivates further investigation of the fracture mechanics and properties of blood clots, and may enable the development of improved hemostatic materials.

## Acknowledgments

This work was supported by New Frontiers in Research Fund – Exploration (grant NFRFE-2018-00751) and Canada Foundation for Innovation (Grant 37719). S.L. and Z.M. acknowledged the financial support from McGill Engineering Doctoral Award. G.B. acknowledged the financial support from National Institute on Deafness and Other Communication Disorders. J.L. acknowledged the support from Canada Research Chair Program.

