## Supplementary information for "Fracture Mechanics of Human Blood Clots: Measurements of Toughness and Critical Length scales"

ARTICLE

Received 00th January 20xx,
Accepted 00th January 20xx

DOI: 10.1039/x0xx00000x

Shiyu Liu[a], Guangyu Bao[a], Zhenwei Ma[a], Christian J. Kastrup[b] and Jianyu Li[ac]

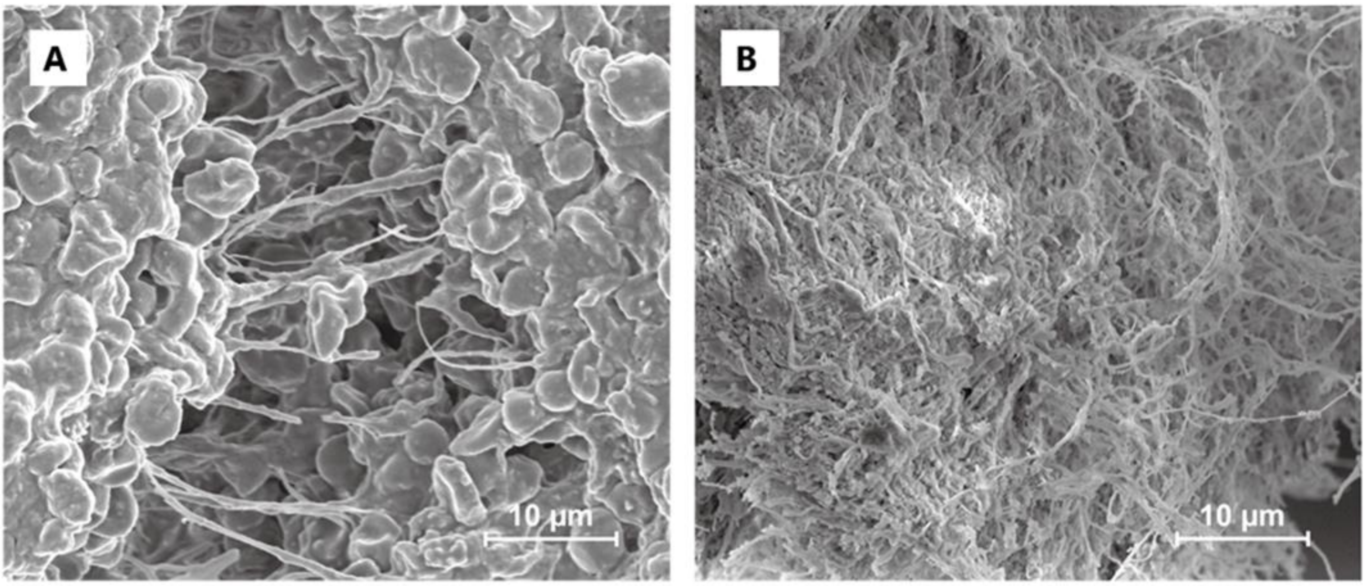


Figure S1 - SEM images of ruptured blood clots. (A) human platelet-poor plasma clot (PPP), (B) human whole blood clot (WB). Scale bar equals 10 µm.

a. *Department of Mechanical Engineering, McGill University, 817 Sherbrooke St W, Montreal, QC H3A 0C3, Canada.*
b. *Michael Smith Laboratories, University of British Columbia, 2185 East Mall, Vancouver, BC V6T 1Z4, Canada.*
c. *Department of Biomedical Engineering, McGill University, 817 Sherbrooke St W, Montreal, QC H3A 0C3, Canada.*
† Footnotes relating to the title and/or authors should appear here.
Electronic Supplementary Information (ESI) available: [details of any supplementary information available should be included here]. See DOI: 10.1039/x0xx00000x

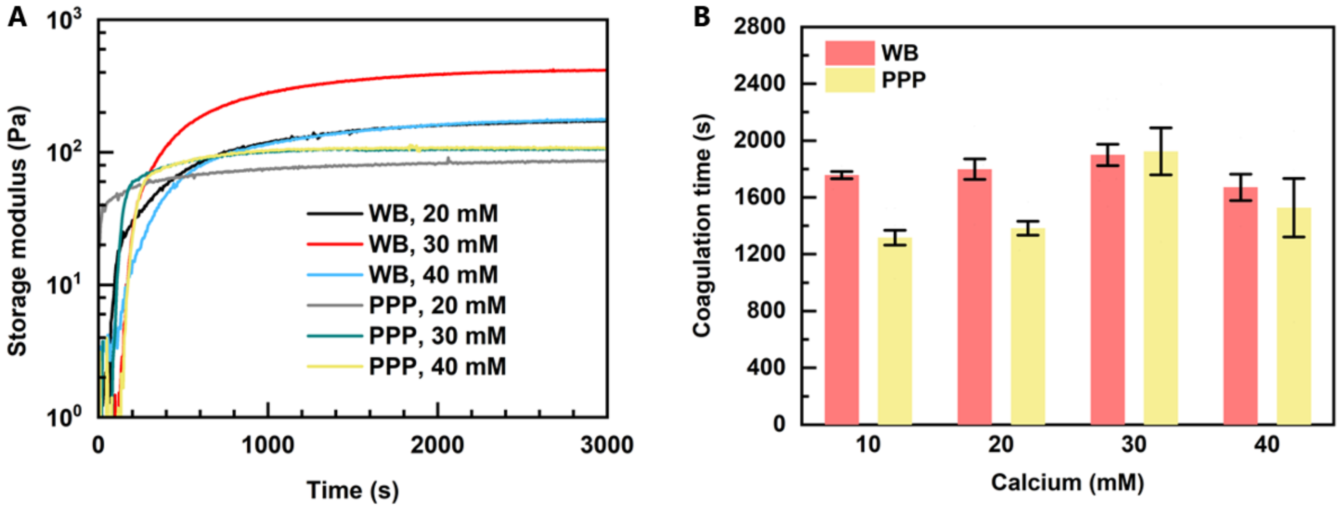


Figure S2 - (A) Gelation kinetics of WB and PPP clots stiffening with varying calcium concentrations. (B) Coagulation time of whole blood and platelet-poor plasma as a function of calcium concentrations. Coagulation time is defined as the time when the storage modulus reaches 90% of plateau value in this study.

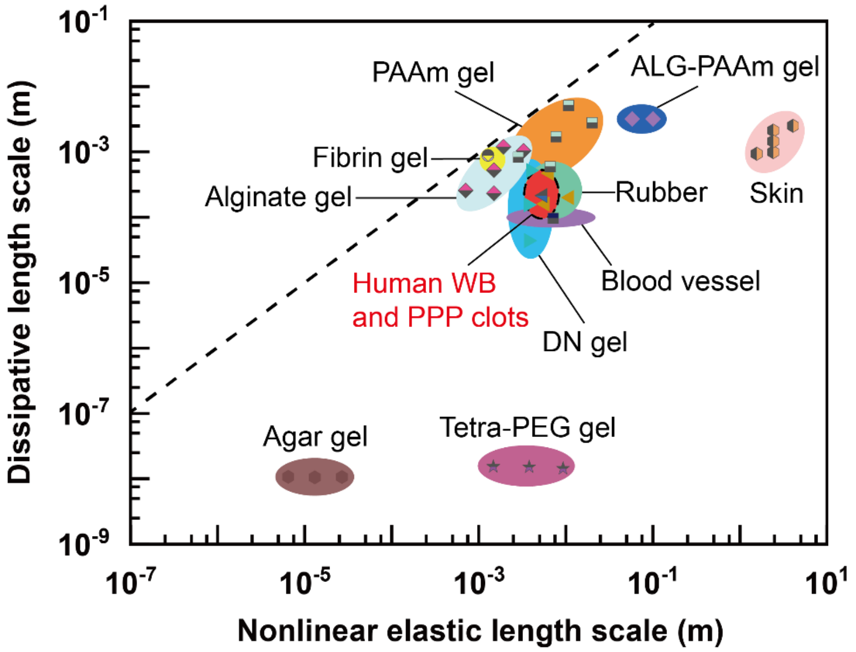


Figure S3 - Fracture length-scale chart with data points for soft and biological materials. Human WB and PPP clots are resulting from this work, while fibrin gel[1], blood vessel[2,3], skin,[4] natural rubber,[5] alginate gels,[4] double-network (DN) gels,[6,7] polyacrylamide (PAAm) gels,[4] alginate-polyacrylamide (ALG-PAAm) gels,[4] agar gels,[8] and Tetra-PEG gels[9] are extracted from the literature. Dash line represents that the material's dissipative length scale and nonlinear length scale are equal.

### Conflicts of interest

The authors declare no known competing financial interests or personal relationships that could have appeared to influence the work reported in this paper.

### Acknowledgements

This work was supported by New Frontiers in Research Fund – Exploration (grant NFRFE-2018-00751) and Canada Foundation for Innovation (Grant 37719). S.L. and Z.M. acknowledged the financial support from McGill Engineering Doctoral Award. G.B. acknowledged the financial support from National Institute on Deafness and Other Communication Disorders. J.L. acknowledged the support from Canada Research Chair Program.

 This journal is © The Royal Society of Chemistry 20xx

Please do not adjust margins




This journal is © The Royal Society of Chemistry 20xx



Please do not adjust margins
